# The role of opioid receptors in tianeptine-induced beta oscillations and memory enhancement

**DOI:** 10.64898/2026.05.10.724133

**Authors:** Margarida J Trigo, Tegan S Knott, Rosamund F Langston, Jeremy J Lambert, Stephen J Martin

## Abstract

Memory impairment is a common and sometimes overlooked feature of major depressive disorder, and cognitive deficits may precede the onset of depressive symptoms in some cases. However, the cognitive benefits of first-line treatments such as SSRIs are mixed. Tianeptine is an atypical antidepressant and cognitive enhancer that neither interacts with monoamine receptors nor inhibits the reuptake of their neurotransmitters. Its antidepressant efficacy in animal models requires activation of the mu-opioid receptor (mu-OR) and phosphorylation of the AMPA receptor. However, the receptors that mediate its memory enhancing actions have never been investigated. We therefore tested the ability of tianeptine to improve spatial memory in a cross-maze task in wild-type (WT) mice compared to its effects in mice with global knockout of either the mu-OR or delta-OR. In parallel, we assessed the effects of tianeptine on hippocampal oscillatory activity and spontaneous locomotion in the same genotypes. Adult male and female WT, mu -/-, and delta -/- mice on a C57BL/6J background were implanted with hippocampal electrodes for the recording of local field potential (LFP) oscillations. Consistent with our previous observations in anaesthetised rats, injection of tianeptine (10 mg/kg and 30 mg/kg SC) caused a dose-dependent increase in beta-frequency power in WT mice that was maximal at circa 25 Hz. The same effect was observed in delta -/- mice, but the increase in beta was completely absent in mu -/- animals. As others have reported previously, tianeptine also caused a mu-OR-dependent increase in spontaneous locomotor activity, but with a time-course that was distinct from the increase in beta power. Separate groups of WT, mu -/-, and delta -/- mice were tested for their ability to learn a food-rewarded spatial memory task in a cross-maze. Over a 20-day training period, sub-groups of each genotype received either tianeptine (10 mg/kg SC) or vehicle injection 30 min before testing. Tianeptine increased the percentage of correct trials and the number of allocentric (place) responses in WT mice, but did not enhance memory in either mu -/- or delta -/- mice, even though both genotypes were able to learn the task. These results indicate that the ability of tianeptine to drive hippocampal beta oscillations is dependent on the mu-OR, whereas its memory-enhancing actions require the presence of both mu- and delta-ORs. The latter result is consistent with the actions of tianeptine on postsynaptic AMPA receptors, and we are currently exploring the signalling pathways involved in this process.

## Introduction

Major depressive disorder (MDD) is a leading worldwide cause of disability and premature mortality (Yan et al., 2024; Chan et al., 2025). Cognitive dysfunction, including deficits in memory and executive function, is a central and often highly distressing symptom that can persist even during periods of remission, especially in cases of melancholic depression (Dillon & Pizzagalli, 2018; Knight et al., 2020; Parker, 2022; Navarra-Ventura et al., 2025; Semkovska et al., 2025). Current first-line treatments such as selective serotonin reuptake inhibitors (SSRIs) have variable efficacy and, when evident, improvements in affective symptoms typically take several weeks to fully develop (Thomas et al., 2013; Boschloo et al., 2023). Results concerning the therapeutic effects of SSRIs on cognitive function are mixed, ranging from no effect to modest improvement (Keefe et al., 2014; Shilyansky et al., 2016; Colwell et al., 2022). For these reasons atypical antidepressants, particularly those that modulate GABAergic or glutamatergic transmission rather than monoaminergic signalling, have attracted considerable clinical interest (Gould et al., 2019; Freudenberg et al., 2025; Fricker et al., 2025). An example of this class of drug is tianeptine, an antidepressant with additional memory-enhancing actions in both humans (Jeon et al., 2014; García-Alberca et al., 2022) and rodents (Jaffard et al., 1991; McEwen et al., 1997; Morris et al., 2001; Zoladz et al., 2008; Zhang et al., 2018).

Tianeptine was first licensed for antidepressant use in France in 1989, and it is currently approved in over 60 countries, though these do not include the UK or USA (Nishio et al., 2024). Although it has a tricyclic chemical structure, tianeptine neither interacts with serotonin or noradrenaline receptors, nor inhibits the reuptake of these neurotransmitters (McEwen et al., 2010). Although this was not recognised at the time of its initial development and licensing, tianeptine is an agonist of µ and δ opioid receptors (µ- and δ-ORs; Gassaway et al., 2014; Singleton et al. 2024), and activation of µ-ORs plays a key role in its antidepressant effects in animal models of depression (Samuels et al., 2017; Han et al., 2022). Also implicated in tianeptine’s antidepressant actions is the drug’s ability to enhance AMPA-receptor-mediated synaptic transmission (Kole et al., 2002; Svenningson et al., 2007; Szegedi et al., 2011; Zhang et al., 2013), an effect that we have recently discovered to be dependent, in area CA1 of the hippocampus, on the activation of δ- and µ-OR receptors (Mariano et al., 2026). Consistent with its enhancement of hippocampal excitatory synaptic transmission, tianeptine also facilitates the induction of long-term potentiation (LTP) of synaptic strength (Shakesby et al., 2002; Rocher at al., 2004; Vouimba et al., 2006; Qi et al., 2009; Szegedi et al., 2011; Zhang et al., 2013 & 2018), and the formation of hippocampus-dependent memory—particularly when memory is initially impaired by stress or other manipulations (Jaffard et al., 1991; McEwen et al., 1997; Morris et al., 2001; Zoladz et al., 2008; Zhang et al., 2018). It also reverses hippocampal shrinkage and dendritic atrophy caused by exposure to chronic stress (Watanabe et al., 1992; Czéh et al., 2001; Liu et al., 2011). However, most investigations of these phenomena predate the discovery of tianeptine’s interactions with opioid receptors. A key goal of the current study was therefore to investigate the role of µ- and δ-ORs receptors in tianeptine’s memory-enhancing actions.

Before studying its role in memory, however, we set out to further characterise the changes in hippocampal network activity induced by tianeptine. We have previously reported that tianeptine causes a disinhibition of hippocampal CA1 pyramidal cells in vivo and in vitro (Martin et al., 2025), an effect that might be mediated by the activation of µ-ORs on inhibitory interneurons such as parvalbumin-positive basket cells (e.g. Shao et al., 2020). Tianeptine also causes naloxone-sensitive beta-frequency oscillations of the local field potential (LFP) in anaesthetised rats (Burt et al., 2026). However, although naloxone’s affinity is highest for the µ-OR, it does not exclude a role for δ-ORs in the generation of beta oscillations. The use of urethane anaesthesia in the earlier study raises additional issues, since urethane potentiates the activation of GABA_A_ receptors, but inhibits NMDA and AMPA receptors (Hara & Harris, 2002). Theta, beta, and gamma-frequency oscillations can still be readily observed under urethane anaesthesia (Martin, 2001), and the architecture of LFP activity resembles that of sleep under these circumstances (Pagliardini et al., 2013). Nevertheless, the frequency, amplitude, and temporal structure of LFP activity are likely to be different under anaesthesia compared to the waking state.

To address these issues, we first studied the effects of tianeptine on hippocampal LFP activity in freely moving mice with chronically implanted electrodes. Using wild-type (WT), δ-OR knockout (δ -/-), or µ-OR knockout (µ -/-) mice, we set out to determine the opioid receptor responsible for tianeptine’s ability to increase beta-frequency oscillations. In parallel, we characterised the dose-dependence, time course, and opioid-receptor-dependence of tianeptine’s locomotor stimulant actions. Lastly, we assessed the memory enhancing effects of tianeptine in a food-rewarded spatial-reference-memory cross-maze task, using δ -/- and µ -/-mice to establish the role of opioid receptors in the pro-cognitive actions of the drug.

## Methods

An overview of experiments and testing schedules is provided in Fig. 1.

**Fig. 1.**
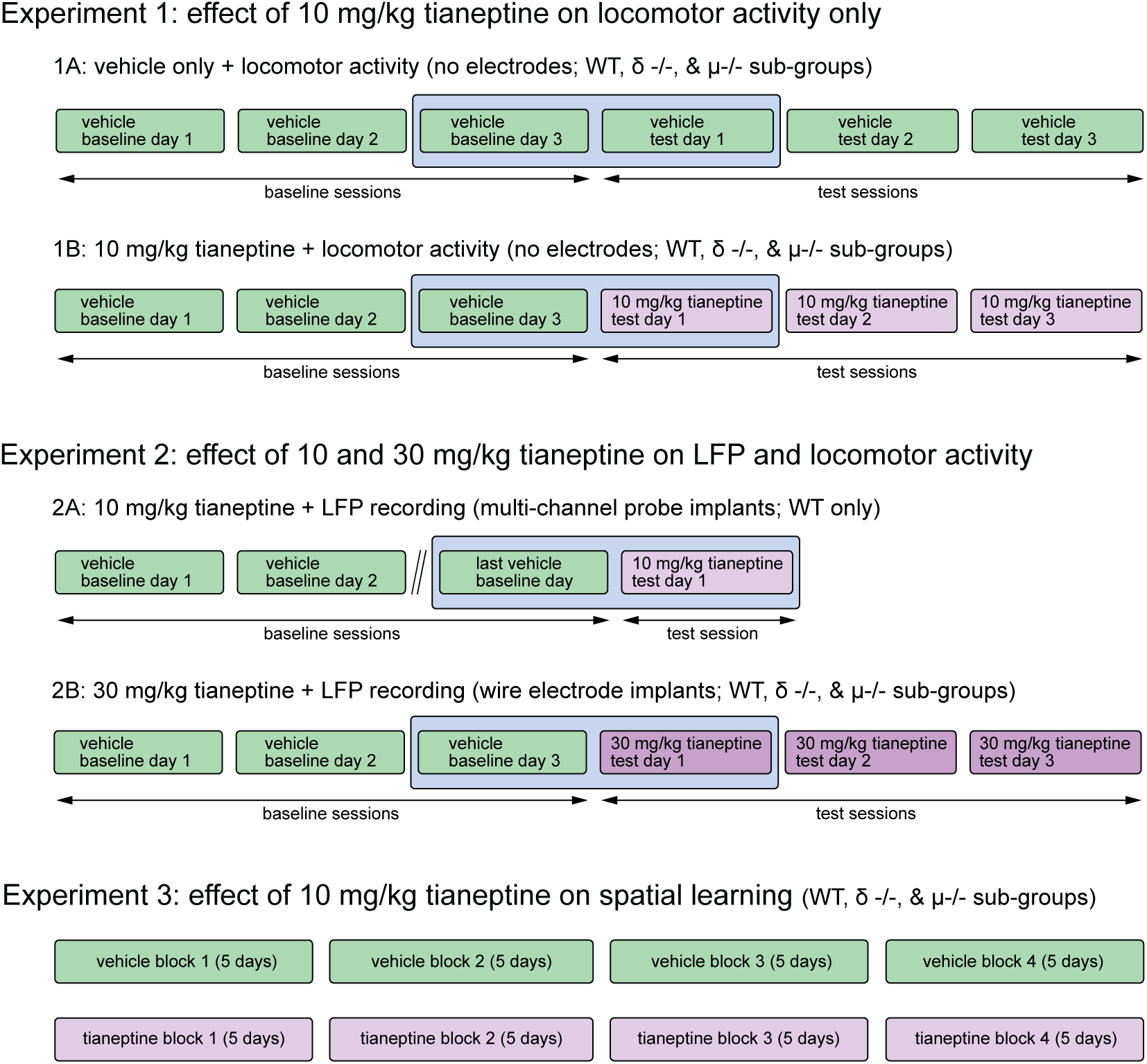
Experimental design and testing schedule. Experiment 1 assessed the effects of vehicle (saline; Experiment 1A) or 10 mg/kg tianeptine (Experiment 1B) on locomotor activity alone, whereas Experiment 2 involved both LFP and locomotor activity monitoring after injection of tianeptine at 10 mg/kg (Experiment 2A) or 30 mg/kg (Experiment 2B). In all cases, the testing schedule comprised a series of days in which drug or vehicle injection was followed by 1-h period of locomotor activity monitoring. This always involved three or more days of habituation to the testing box after vehicle injection (baseline sessions), followed by one or more days of testing (test sessions) after either continued vehicle (Experiment 1A) or tianeptine (Experiments 1B, 2A, & 2B) administration. In most cases, the schedule comprised exactly 3 days of baseline monitoring followed by 3 test sessions in WT, δ -/-, and µ -/- mice (Experiments 1A, 1B, & 2B). The sole exception was Experiment 2A in which WT mice alone were tested; they received 3-6 days of baseline monitoring after vehicle injection, followed by a single day of 10 mg/kg tianeptine administration. In all cases, however, the key comparison was the within-subjects increase in activity or LFP power between the final baseline session and first test session, indicated by the pale blue box in which these two days are highlighted. Experiment 3 was a purely behavioural study of the performance of WT, δ -/-, and µ -/- mice in a cross-maze spatial memory task. Training comprised four 5-day blocks of 8 trials each. In this case, the key comparison was a between-subjects analysis of the difference in performance between vehicle and 10-mg/kg-tianeptine-treated sub-groups.

### Animals

All experiments were carried out under the authority of licences granted by the UK Home Office under the Animals (Scientific Procedures) Act 1986. Prior to submission to the Home Office, applications for project licences were approved by the University Welfare and Ethical Use of Animals Committee, acting in its capacity as an Animal Welfare and Ethical Review Body as required under the Act.

Unoperated animals and mice undergoing behavioural testing were group-housed, but mice implanted with electrodes for LFP recording were singly housed after surgery. Access to water and food (standard rodent pellets supplemented with wheat and sunflower grains) was unrestricted in all cases except for those mice undergoing food restriction during the main testing phase of the cross-maze study (see below). Mice were maintained on a 12-h light-dark cycle at a constant temperature (19-24 °C). Cages contained sawdust with shredded cardboard, cotton-fibre nesting material, and a cardboard ‘house’. Enrichment items included wooden chew-sticks, a red polycarbonate tunnel, a cable-tie ‘swing’, and wheat and sunflower grains. Ages ranged from 2-3 months at the start of testing. Male and female wild-type (WT) C57Bl/6J mice were obtained from Charles River Laboratories (Margate, UK). Knockout mice were bred in-house; µ -/- mice were originally generated by Matthes et al. (1996) (B6.129S2-*Oprm1^tm1Kff^*/J; Jackson Laboratory strain #007559) and δ -/- mice were created by Filliol et al. (2000) (B6.129S2-*Oprd1^tm1Kff^*/J; Jackson Laboratory strain #007557). Both lines were fully backcrossed onto a C57Bl/6J background.

### Drug solutions

Tianeptine sodium (Kemprotec Limited, Cumbria UK) was dissolved in sterile injectable saline (vehicle) at a concentration of 2 mg/ml or 6 mg/ml. Injections were administered subcutaneously (SC) at 5 ml/kg, corresponding to tianeptine doses of 10 mg/kg or 30 mg/kg.

### Locomotor testing

Locomotor activity monitoring was carried out in clear acrylic cubes with dimensions of 30 x 30 x 30 cm, and open at the top. Behaviour was monitored via overhead cameras connected to a PC running Any-maze software (Stoelting Europe, Dublin, Ireland). For mice undergoing locomotor activity monitoring without LFP recording (Experiment 1 in Fig. 1), testing was carried out in a series of 4 boxes, separated by opaque barriers. Testing always comprised a 1-h session; injections occurred immediately before the mouse was placed in the activity box. Testing began with a 3-day period of locomotor testing after vehicle injection designed to habituate the mice to the apparatus and achieve a stable level of exploratory activity across days. This was followed by a further 3-day period during which mice received either further vehicle injections (Experiment 1A in Fig. 1) or 10 mg/kg tianeptine injection (Experiment 1B in Fig. 1). For mice undergoing tethered LFP recording at the same time as locomotor monitoring (Experiment 2 in Fig. 1), testing was carried out in identical acrylic boxes located within wooden recording chambers. Further details, including the testing schedule, are given below.

### LFP recording

#### Surgery

Mice were implanted with electrodes for LFP recording under isoflurane anaesthesia. They were placed in a stereotaxic frame fitted with an anaesthetic nose-cone, and body temperature was maintained using a rectal thermometer and heat pad connected to a homeothermic monitoring unit (Harvard Apparatus, Holliston, MA, USA). Carprofen (5mg/kg SC) was given for analgesia at induction and postoperatively, lidocaine was applied to the ear canal and scalp before ear-bar insertion, and a lubricating gel was applied to the eyes to prevent corneal drying. Under aseptic conditions, an incision was made in the skin and periosteum and a small burr-hole was made at the target location. The electrode implanted was either a 16-channel single-shank probe (NeuroNexus A1x16-3mm-100-703-CM16; Ann Arbor, MI, USA) (Experiment 2A in Fig. 1) or a conventional wire electrode comprising two twisted insulated platinum / iridium wires (diameter = 0.075 mm; Advent Research Materials, Eynsham, UK), connected to an 18-pin wire adapter (#B7600; Intan Technologies, Los Angeles, CA, USA) (Experiment 2B in Fig. 1). These were lowered into the hippocampus; coordinates relative to bregma were AP = −2.0 mm; ML = 2.0 mm; depth relative to the dura = −2.1 mm for multi-channel probes and −1.3 mm for wire electrodes. Small jewellers’ screws were also secured to the skull to serve as anchor point, one of which, positioned in the occipital bone, served as a reference and ground. The whole assembly was then secured to the skull using dental cement. Mice were placed in an incubator and monitored during recovery from anaesthesia. Testing did not begin for a period of at least one week after surgery, and after mice had fully recovered based on their appearance, weight, and behaviour.

#### LFP data acquisition and testing procedure

Recording was carried out in a 30 x 30 x 30 cm open-topped acrylic cube located inside a fan-ventilated wooden recording chamber with dimensions of circa 50 x 50 x 50 cm, and fitted with ceiling lights to provide diffuse lighting. The inside walls of the cabinet were electrically shielded using grounded aluminium foil coated with adhesive white plastic. A video camera and commutator assembly were secured to the roof of the chamber, the latter comprising a 12-channel Adafruit slip-ring (product #1196; Adafruit Industries, New York City, USA) mounted in a 3-D printed assembly (Patrick Spooner, University of Edinburgh) and fitted with two Intan 18-pin electrode adapter boards (product #C3418; Intan Technologies, Los Angeles, USA). A spring, 10-cm in diameter, prevented the cable from becoming slack, and transmitted small rotational forces to the commutator. The output adapter board was connected via Intan RHD SPI interface cables (product #C3203) to an OpenEphys acquisition system (OpenEphys, Atlanta, GA, USA), and the input board was connected, via the same cable type, to an Intan 16-channel headstage (product #C3335) that was plugged into the electrode implant during testing. The wide-band LFP signal was sampled at 30 kHz and data from all channels was recorded via the OpenEphys system.

Mice were habituated to the recording chamber and the connection of a headstage and cable over a 3-day period before the start of data collection. On the first day, they were placed inside the recording chamber for 15 min with lights and fan on, but the recording cable was not attached. The second day followed the same procedure except that the cable and headstage were connected, and this was repeated for a 45-min period on the third day. The main phase of testing and data collection started after this. Testing began in all cases with a 3-day period during which mice were injected with vehicle (sterile saline) immediately before a 1-h recording session. For mice with multi-channel probes, this was followed for half the mice by a single recording session after 10 mg/kg tianeptine injection; the other half underwent 3 additional days of recording after vehicle injection, followed by a single recording session after 10 mg/kg tianeptine injection (Experiment 2A in Fig. 1). For mice implanted with wire electrodes, the initial 3-day period of recording after vehicle injection was followed by a 3-day period of recording after 30 mg/kg tianeptine injection (Experiment 2B in Fig. 1). In all cases, the analysis of LFP data focussed on the contrast between baseline activity during the final day of vehicle administration, and LFP activity during the first day of tianeptine administration.

#### LFP analysis

For both wire electrodes and multi-channel probes, the channel with the highest baseline spectral power in the theta-frequency range was selected for further analysis; in the latter case, this channel was identified using a custom-written Python script written by Christian Keitel, University of Dundee. Data were then imported into Brainstorm (Tadel et al. 2011), which is documented and freely available for download online under the GNU general public license (http://neuroimage.usc.edu/brainstorm). Data were low-pass filtered at 500 Hz, and resampled at 1000 Hz. Spectral analysis was performed on successive 2-s samples of data (50% window overlap) using Welch’s power spectral density estimate, and mean spectral power over the 0-90 Hz range was exported in 2-min bins. Time-frequency heat maps were constructed by plotting 2-min power spectra as a 3-D time-frequency graph with power as the z-axis. The change in spectral power between treatment and baseline days was calculated by subtracting the log_10_ power at each time point and frequency bin on a treatment day from the corresponding time point and frequency bin on the baseline day. These data were also plotted as a 3-D time-frequency heat map, this time with change in spectral power relative to baseline as the z-axis. The mean change in spectral power in the beta-frequency band, defined as 15-30 Hz, was also calculated.

### Cross-maze spatial learning

The cross-maze task was adapted from the procedure described by Le Merrer et al., 2013. It was made from clear acrylic and comprised 4 arms (35 cm long, 8 cm wide, and 33 cm high) connected to a square central section (8 cm x 8 cm). Four removable sliding gates (7.5 cm wide and 33 cm high) located 10 cm from the closed end of each arm formed ‘start boxes’. Four identical gates were located at the junction between each arm and the central section. The ends of the east (E) and west (W) arms featured holes for the insertion of a ball-bearing sipper tube connected to a feeding bottle. The cross-maze was located in the centre of a room containing prominent extra-maze landmarks such as a blue dividing panel, posters, and laboratory furniture. An overhead video camera was connected to a PC running Any-maze software for behavioural analysis.

All mice were handled by the experimenter for 5 days before the start of testing. During this period, a feeding bottle containing soya milk was made available in the home cage for 30 min per day over 3 consecutive days. Mice were habituated to the maze during three 10-min sessions during which they were allowed to explore the apparatus. At the start of each session, they were confined to a start box for 15 s before being released. During the first habituation session, the mice were allowed unrestricted exploration. In the second and third sessions, upon their first entry into the E or W arms, mice were confined in the chosen arm for 20 s by closure of the gate separating the arm from the central compartment. The gate was then re-opened, and mouse were allowed unrestricted exploration for the remainder of the session. During the third habituation session, soya milk reward was available at the ends of the E and W arms. Food restriction commenced after cross-maze habituation, and 48 h before the start of the main phase of cross-maze testing. Mice were weighed daily and food was adjusted to maintain a bodyweight no less than 85% of the free-feeding value.

The main phase of cross-maze testing was carried out over 20 days, divided into weekly blocks of 5 days each. Mice were assigned randomly to counterbalanced combinations of start locations (N or S for training trials) and reward locations (E or W). During training trials, soya milk was only available from the bottle located in the assigned goal arm. The bottle in the opposite arm contained tissue paper soaked in soya milk, but none was available via the sipper, a procedure designed to thwart strategies based on odour cues rather than memory. At the start of each training trial, a mouse was confined to either the north (N) or south (S) start box for 15 s before the gate was lifted. The arm opposite to the start box was always closed. As soon as a mouse entered either the E or W arm, the gate separating the chosen arm from the central area was closed, and the mouse was confined to the arm for 20 s before being removed from the maze and placed in a holding box for 2 min before the next trial began. Latency to choose an arm was calculated using Any-maze as the time between an animal’s release from the start box, until its whole body entered either the E or W arm.

Probe trials were conducted at the start of every testing session, before the 7 training trials. On probe trials, mice were placed in the start box opposite to the location assigned for training trials. Soya milk reward was available from both E and W arms during these trials since their purpose was to assess the strategy used, not to reward or punish egocentric versus allocentric responding. Egocentric responding during probes was defined as a response involving a turn in the same direction (i.e. left or right) as would lead to the reward arm during training trials, but leading to the opposite cardinal location. An allocentric response is conventionally defined as one that leads to the same cardinal location, but requires an opposite direction of turning to that required during training trials. In this experiment, a more stringent measure of allocentric responding was defined by an allocentric response during a probe, followed by a correct response on the subsequent training trial. This measure requires a mouse to make turns in opposite egocentric directions across two consecutive trials in order to reach the same allocentric location on both occasions.

## Results

### Effect of tianeptine on locomotor activity

Tianeptine caused a dose-dependent increase in locomotor activity, a summary of which is presented in Fig. 2. This analysis draws on data from across the series of experiments outlined in Fig. 1. We compared data from Experiments 1A (vehicle only + locomotor activity), 1B (10 mg/kg tianeptine + locomotor activity), and 2B (30 mg/kg tianeptine + LFP recording), each of which comprised three days of vehicle injection (‘vehicle baseline’ days), followed by three test days during which vehicle, 10 mg/kg tianeptine, or 30 mg/kg was administered (Fig. 2A). In all of these experiments, separate sub-groups of WT, δ -/-, and µ -/- mice were tested. In Experiments 1A and 1B, animals were untethered and no electrodes were implanted. In contrast, in Experiment 2B, the ‘30 mg/kg tianeptine + LFP recording’ group had chronically implanted electrodes, and locomotor activity was monitored at the same time as LFP recording. Comparisons of locomotor activity in the latter versus the former groups should therefore be treated with caution owing to the presence of a cable for tethered recording and other differences in the testing procedure and environment (see Materials and Methods). Nonetheless, we present evidence below that these differences had minimal impact on locomotor activity.

**Fig. 2.**
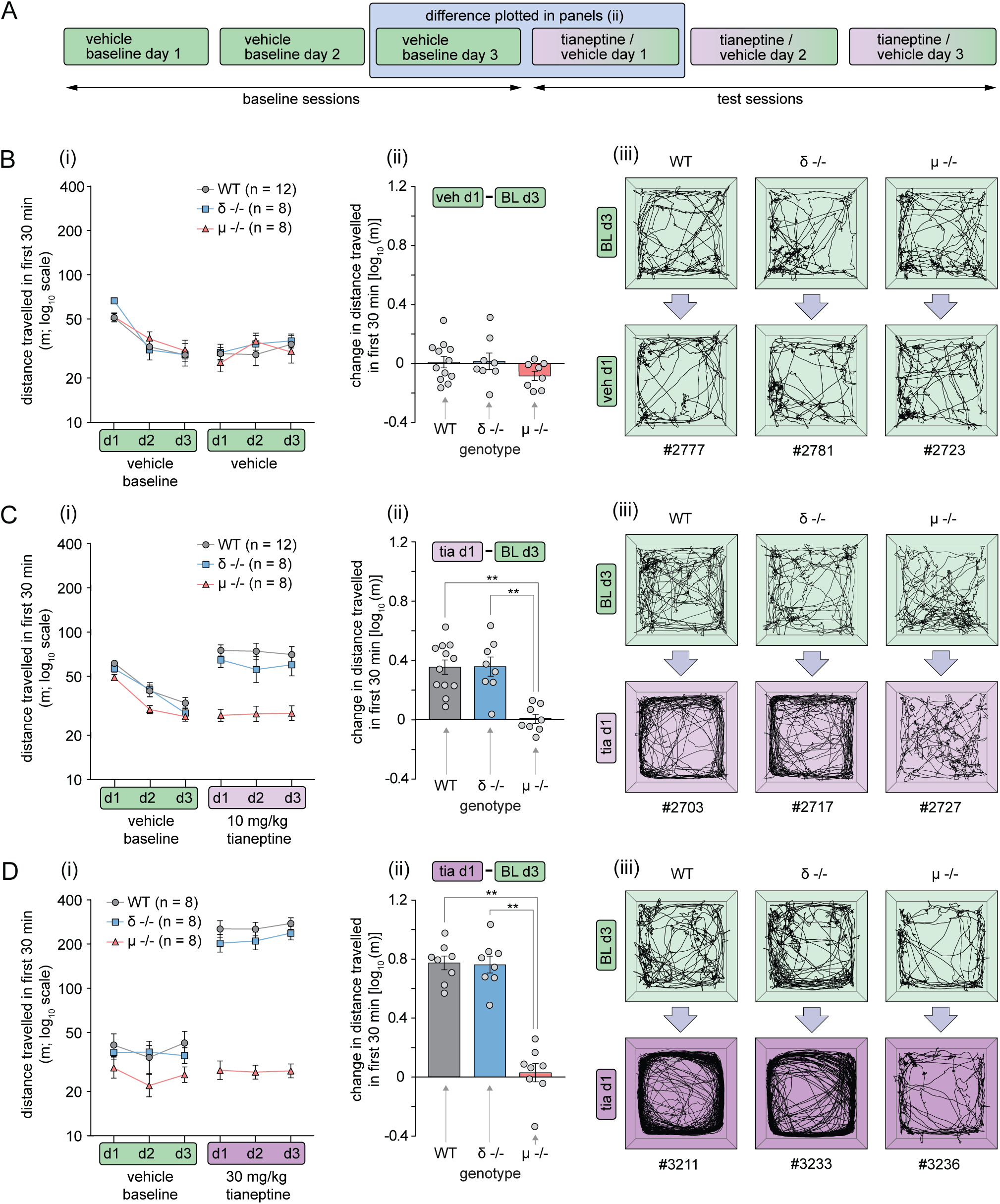
Summary of tianeptine’s acute dose-dependent locomotor effects. (A) Experimental timeline illustrating the three baseline sessions (vehicle injection) followed by three test sessions (vehicle, 10 mg/kg, or 30 mg/kg tianeptine injection). (B) Locomotor activity in control mice injected with vehicle throughout testing (Experiment 1A in Fig. 1): (i) Distance travelled (log_10_ scale) during baseline and test sessions in WT, δ -/-, and µ -/- mice; (ii) change in distance travelled [units = log10 (m)] between vehicle baseline day 3 (BL d3) and vehicle test day 1(veh d1); (iii) Representative examples of the paths taken by individual mice of each genotype over the first 10 min of vehicle baseline day 3 and vehicle test day 1. There were no changes in the level or pattern of activity across days. (C) Locomotor activity in mice injected with 10 mg/kg tianeptine during test sessions (Experiment 1B in Fig. 1): (i) Distance travelled (log_10_ scale) during baseline and test sessions in WT, δ -/-, and µ -/- mice; (ii) change in distance travelled [units = log10 (m)] between vehicle baseline day 3 (BL d3) and vehicle test day 1(veh d1). **p < 0.01; post-hoc pairwise comparisons (Tukey’s HSD; see main text). (iii) Representative examples of the paths taken by individual mice of each genotype over the first 10 min of vehicle baseline day 3 and vehicle test day 1. Note the increase in activity in WT and δ -/- animals, but not in µ -/- mice. (D) Locomotor activity in mice injected with 30 mg/kg during test sessions (Experiment 2B in Fig. 1): (i) Distance travelled (log_10_ scale) during baseline and test sessions in WT, δ -/-, and µ -/- mice; (ii) change in distance travelled [units = log10 (m)] between vehicle baseline day 3 (BL d3) and vehicle test day 1(veh d1). **p < 0.01; post-hoc pairwise comparisons (Tukey’s HSD; see main text). (iii) Representative examples of the paths taken by individual mice of each genotype over the first 10 min of vehicle baseline day 3 and vehicle test day 1. Note the striking increase in activity in WT and δ -/- animals, an effect that was absent in µ -/- mice.

Fig. 2B(i) shows distance travelled in the ‘vehicle only + locomotor activity group’ (Experiment 1A in Fig. 1) over the first 30 min of each session across the 3 vehicle baseline days (left-hand side) followed by the 3 vehicle test days (right-hand side). Significant deviations from homogeneity of variance were frequently observed in the distance travelled measure. For this reason, data were plotted on a log_10_ scale, and log_10_-transformed for statistical analysis. In this case, distance travelled decreased significantly across the 3 vehicle baseline days [F(2,50) = 69.7; p < 0.001] indicating habituation to the apparatus, with no difference between genotypes [F(2,25) = 0.22; p = 0.80]. There was also a significant change in distance travelled across the three vehicle test days [Fig. 2B(i); right-hand side] owing to a slight overall rise in activity [F(2,50) = 4.13; p = 0.022], but again no difference between genotypes [F(2,25) = 0.15; p = 0.86]. Analysis of the change in distance travelled between vehicle baseline day 3 and vehicle test day 1 [Fig. 2B(ii)] did not reveal any significant changes [p > 0.09 in all cases; one-sample t-tests (comparisons to zero) with Bonferroni correction], and there was no overall effect of genotype [F(2,25) = 1.51; p = 0.24]. Representative examples of the paths taken by one mouse from each genotype over the first 10 min of vehicle baseline day 3 (upper panels) and vehicle test day 1 (lower panels) are shown in Fig. 2B(iii). The paths indicate a typical pattern of low-intensity exploratory activity that did not change across days. This is expected because animals were habituated to the apparatus at this point, yielding stable activity levels across days, and mice were injected with vehicle at the start of each session.

Fig. 2C shows locomotor activity data from the ‘10 mg/kg tianeptine + locomotor activity’ group [Experiment 1B in Fig. 1]. Distance travelled again decreased across the 3 baseline vehicle days [F(2,50) = 76.4; p < 0.001; Fig. 2C(i); left-hand side], and on this occasion there was also a significant effect of genotype [F(2,25) = 4.42; p = 0.023], with significantly lower baseline activity in the µ -/- group compared to WTs (p = 0.018; post-hoc Tukey’s HSD). There was no overall change in activity across the three vehicle test days [F(2,50) = 0.26; p = 0.78; Fig. 2C(i); right-hand side], but a highly significant main effect of genotype [F(4,50) = 20.2; p < 0.001], with significantly higher activity in the WT and δ -/- groups relative to the µ -/- group (p < 0.001 in both cases; post-hoc pairwise comparisons; Tukey’s HSD). Analysis of the change in distance travelled between vehicle baseline day 3 and vehicle test day 1 [Fig. 2C(ii)] revealed a significant effect of group [F(2,25) = 14.6; p < 0.001], with significantly increased locomotor activity in the WT and δ -/- groups relative to the µ -/- group (p < 0.001 in both cases; post-hoc pairwise comparisons; Tukey’s HSD). This increase was highly significant in WT and δ -/- groups (p < 0.001 in both cases; one-sample t-tests with Bonferroni correction). Representative examples of the paths taken by one mouse from each genotype over the first 10 min of vehicle baseline day 3 (upper panels) and vehicle test day 1 (lower panels) are shown in Fig. 2C(iii). Note the increase in locomotor activity in WT and δ -/- mice, but the absence of any change in µ -/-animals.

Fig. 2D shows locomotor activity data from the ‘30 mg/kg tianeptine + LFP recording’ group [Experiment 2B in Fig. 1]. In this group, there was no significant decrease in distance travelled over the three baseline vehicle days [F(2,42) = 1.18; p = 0.32; Fig. 2D(i); left-hand side], likely because these mice had previous experience of the testing apparatus during habituation to the presence of the recording cable (see Materials and Methods for details). To confirm that these and other procedural differences had minimal effects on baseline locomotor activity by vehicle baseline day 3, we conducted a two-way ANOVA of activity on this day with genotype and experiment type [locomotor testing only (Experiments 1A & 1B) versus locomotor testing + LFP recording (Experiment 2B)]. Note that this analysis focuses on baseline activity only, since tianeptine was not administered on this day. The ANOVA did not reveal a significant overall effect of experiment type [F(1,74) = 2.82; p = 0.097] despite the large number of animals included in this analysis (total n= 80). This confirms that the presence of a recording cable, and other minor differences in experimental procedure, had little effect on the key baseline locomotor measure used in subsequent analysis. In the ‘30 mg/kg tianeptine + LFP recording’ group, there was also no significant overall effect of genotype on vehicle baseline days [F(2,21) = 2.09; p = 0.15; Fig. 2D(i); left-hand side]. Over the 3 tianeptine test days [Fig. 2D(i); right-hand side], there was no overall change in activity [F(2,42) = 1.36; p = 0.27], but a significant effect of genotype [F(2,21) = 134.4; p < 0.001]. Post-hoc pairwise comparisons (Tukey’s HSD) revealed significantly higher activity in the WT and δ -/- groups relative to the µ -/- group (p < 0.001 in both cases). Analysis of the change in distance travelled between vehicle baseline day 3 and vehicle test day 1 [Fig. 2D(ii)] revealed a significant effect of genotype [F(2,21) = 58.5; p < 0.001], with significantly higher locomotor activity in the WT and δ -/- groups relative to the µ -/- group (p < 0.001 in both cases; post-hoc pairwise comparisons; Tukey’s HSD). This increase was highly significant in WT and δ -/- groups (p < 0.001 in both cases; one-sample t-tests with Bonferroni correction). Representative examples of the paths taken by one mouse from each genotype over the first 10 min of vehicle baseline day 3 (upper panels) and vehicle test day 1 (lower panels) are shown in Fig. 2D(iii). Note the marked increase in locomotor activity in WT and δ -/-mice, but the absence of any change in µ -/- animals.

An omnibus ANOVA of the increase in locomotor activity between vehicle baseline d3 and test day 1 including all tianeptine doses [0 mg/kg, i.e. vehicle [Fig. 2B(ii)], 10 mg/kg [Fig. 2C(ii)], and 30 mg/kg [Fig. 2D(ii)] revealed a highly significant increase in activity with increasing dose [F(2,71) = 115.2; p < 0.001], and a significant dose x genotype interaction [F(4,71) = 20.2; p < 0.001]. Analysis of simple main effects revealed a significantly larger increase in activity with increasing tianeptine doses in WT [F(2,71) = 63.9; p < 0.001] and δ -/- mice [F(2,71) = 93.7; p < 0.001], but no effect of increasing dose in µ -/- mice [F(2,71) = 1.32; p = 0.27].

Having established the dose-dependence and requirement for the µ-OR in tianeptine’s locomotor actions, we next focussed on the time course of tianeptine’s actions in unoperated mice (Fig. 3A & 3B-D; see Experiment 1B in Fig. 1), and mice undergoing LFP recording (Fig. 3A & 3E-H; see Experiment 2A & 2B in Fig. 1). For clarity, and because the focus here is on within-subjects changes, rather than between-subjects analyses of groups with heterogeneous variance as in Fig. 2, data are plotted on linear rather than log_10_ scales. Fig. 3B(i) shows the effects of injection of 10 mg/kg on the time course of locomotor activity alongside data from the preceding vehicle baseline day. The difference in activity between these two days is plotted in Fig. 3B(ii). Time points at which activity was significantly increased (one-sample t-tests with Bonferroni correction) are indicated in the figure; an increase in activity was observed all time-points within the first 10 min after injection, except during the very first 2 min. The increase in locomotion was similar in magnitude and time course in δ -/- mice (Fig. 3C). One-sample t-tests with Bonferroni correction, revealed significant increases at equivalent time points to those affected in WT mice [Fig. 3C(ii)]. No increase in locomotor activity was seen in µ-OR -/- mice (Fig. 3D).

**Fig. 3.**
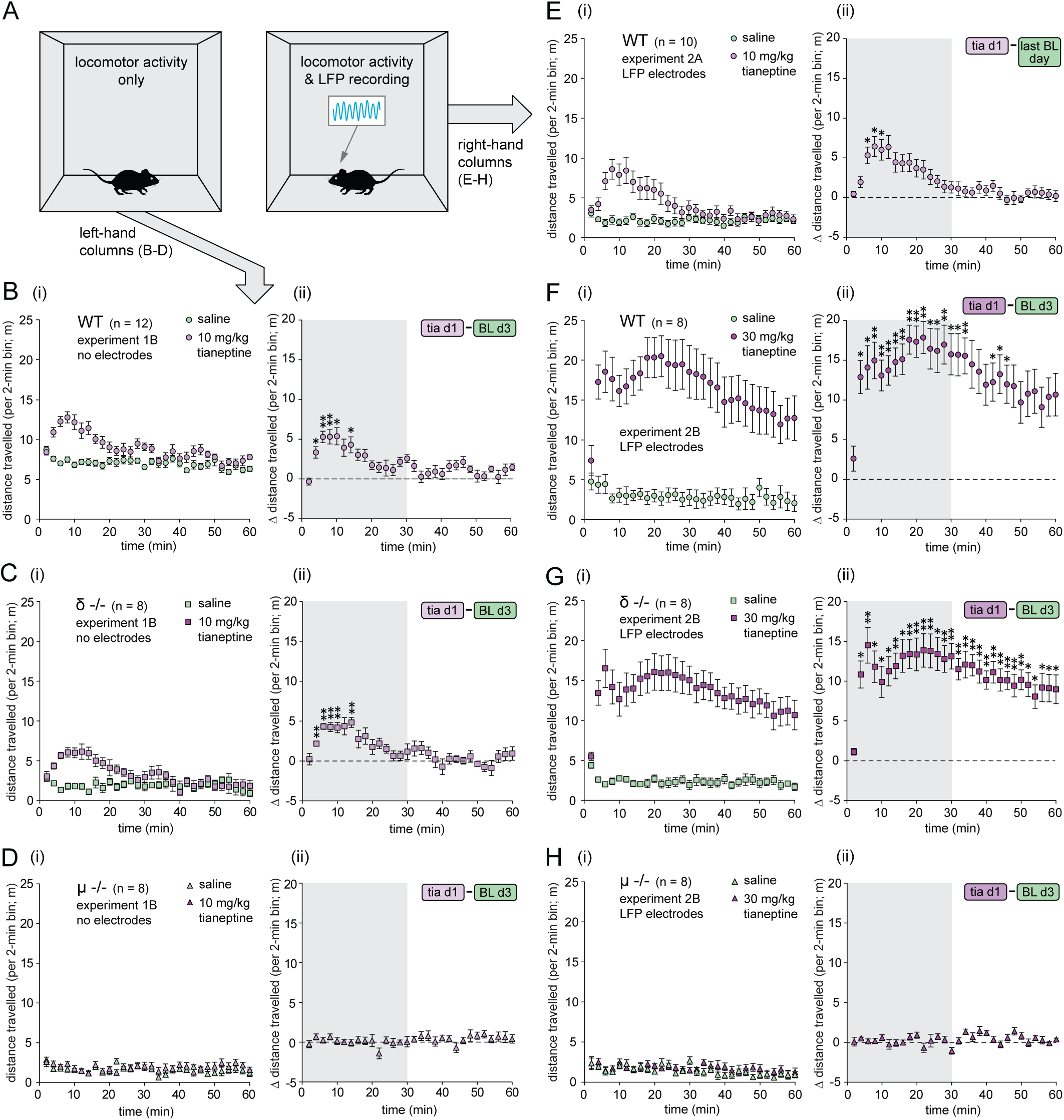
Time-course of tianeptine’s acute locomotor effects. (A) Experimental conditions: the left-hand box illustrates a mouse undergoing locomotor activity monitoring only, corresponding to the left-hand panels B-D (Experiment 1B in Fig. 1), whereas the right-hand box shows a mouse undergoing combined LFP and locomotor activity monitoring, corresponding to the right-hand panels E-H (panel E: Experiment 2A; panel F-H: Experiment 2B; see Fig. 1). (B)(i) Distance travelled by WT mice without electrode implants. Data are plotted in 2-min time bins over the course of the last vehicle baseline day and the first day on which 10 mg/kg tianeptine was administered. (B)(ii) Time course of the increase in distance travelled [first 10 mg/kg tianeptine day (tia d1) – vehicle baseline day 3 (BL d3)]. In subsequent panels, corresponding data are plotted for δ -/- mice without electrode implants injected with 10 mg/kg tianeptine (C), µ -/-mice without electrode implants injected with 10 mg/kg tianeptine (D), WT mice with multi-channel probe implants injected with 10 mg/kg tianeptine (E), WT mice with wire electrode implants injected with 30 mg/kg tianeptine (F), δ -/- mice with wire electrode implants injected with 30 mg/kg tianeptine (G), and µ -/- mice with wire electrode implants injected with 30 mg/kg tianeptine (H). In panels ii: *p < 0.05; ⁑p < 0.001; one-sample t-tests with Bonferroni correction.

Analysis of the locomotor increase in mice undergoing LFP recording revealed a similar increase in activity in WT mice after 10 mg/kg tianeptine injection (Fig. 3E; Experiment 2A in Fig 1). One-sample t-tests with Bonferroni corrections, revealed a significant increase in activity at all time-points within the first 10 min after injection, except during the first two 2-min time-points [Fig. 3E(ii)]. A comparison of the change in activity in mice with and without LFP electrodes indicated no difference between the two groups [Fig. 3E(ii) versus Fig. 3B(i): F(1,20) = 0.022; p = 0.883; ANOVA]; the same was true of baseline activity in the two groups [Fig. 3E(i), vehicle versus Fig. 3B(ii), vehicle: F(1,20) = 0.18; p = 0.68]. This is consistent with the analysis of baseline activity in Fig. 2, and confirms that despite the differences in recording environments, surgical history, and the presence or absence of a cable for tethered recording, locomotor activity was minimally affected in mice undergoing LFP recording relative to unoperated controls.

Fig. 3F shows the locomotor response to 30 mg/kg tianeptine in WT mice undergoing LFP recording. The increase was more pronounced and longer in duration than that observed after injection of 10 mg/kg. The peak increase occurred >20 min after injection, and activity remained elevated throughout the recording session. Activity was significantly increased at all time-points within the first 30 min after injection, except during the first 2 min [Fig. 3F(ii)]. A similar increase in activity was seen at this dose in δ -/- mice (Fig. 3G). Activity was significantly increased at all time points except the first 2 min [Fig. 3G(ii)]. No change in activity was evident in µ -/- mice (Fig. 3H).

### Effect of tianeptine on LFP activity

Tianeptine caused a dose-dependent increase in beta-frequency hippocampal LFP activity (Fig. 4). Like the analysis of locomotor activity above, our analysis focusses on the contrast between the last baseline day preceding tianeptine administration, and the first day on which tianeptine was administered. Fig. 4A(i) shows a mean heat map of spectral power (z-axis) plotted a function of both frequency and time during the last vehicle baseline day recorded in mice implanted with multi-channel probes (Experiment 2A in Fig. 1). For each mouse, the single channel with the highest theta power was selected for spectral analysis, a procedure that ensures that all recordings originate from a similar location in the distal apical dendrites of CA1, close to the hippocampal fissure; in almost all cases, this was also the channel with the highest baseline beta-frequency power (see Materials and Methods). Note the prominent peak at ∼8 Hz, corresponding to hippocampal theta activity. After 10 mg/kg tianeptine injection, a modest increase in beta-frequency power is evident within the first few min after injection [Fig. 4A(ii)]. Fig. 4A(iii) summarises this change in a plot of spectral power versus frequency at the 6-8 min time period over which the increase in beta power was maximal. Injection of the higher dose of 30 mg/kg tianeptine produced a larger increase in beta power in mice in mice implanted with conventional wire electrodes (Fig. 4B) (Experiment 2B in Fig. 1). In this case, the peak increase in beta power occurred 4-6 min after injections [Fig. 4B(ii)], and this is the time period represented in Fig. 4B(iii). A similar increase in beta power was observed in δ -/- mice (Fig. 4C), but no change in LFP activity was evident in µ -/- mice (Fig. 4D).

**Fig. 4.**
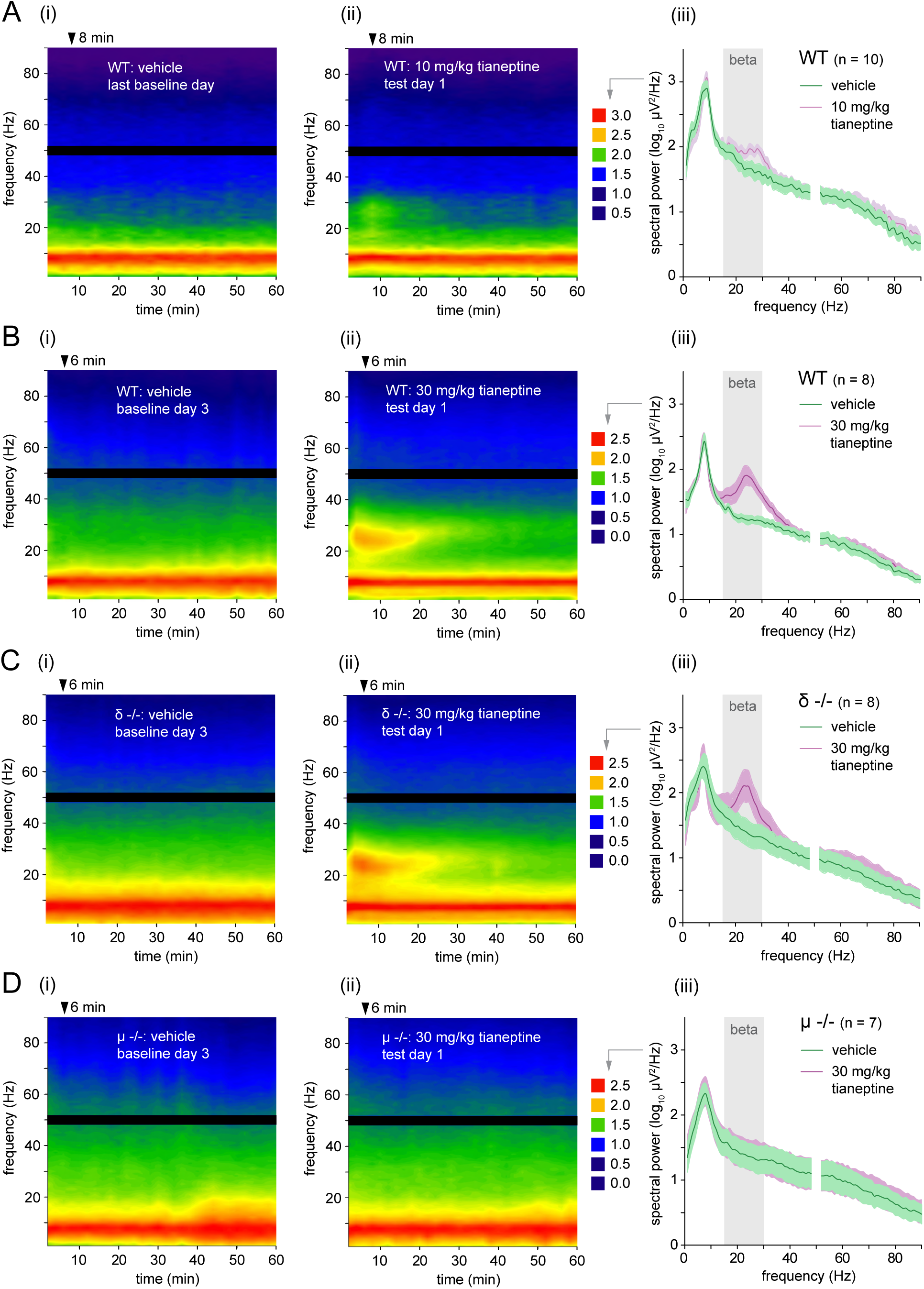
Spectral power before and after tianeptine administration. (A)(i) Heat map of spectral power (z-axis) on the last vehicle baseline day in WT mice implanted with multi-channel probes; data are presented from the channel with the highest power in the theta-frequency band. Power is plotted as a function of frequency (y-axis) and time after vehicle injection (x-axis) (A)(ii) Corresponding heat map after 10 mg/kg tianeptine injection (Experiment 2A in Fig. 1). The colour-coded key indicates log_10_ spectral power (z-axis) in both panels i and ii. (A)(iii) Mean spectral power as a function of frequency over the 2-min period, 6-8 min after injection, during which the increase in power in the beta-frequency range (15-30 Hz; grey bar) was maximal after 10 mg/kg tianeptine administration [see Fig. 5B(iii)]. Subsequent panels show data plotted in the same way for mice with wire electrode implants injected with 30 mg/kg tianeptine (experiment 2B in Fig. 1)—WT (B), δ -/- (C), and µ -/- (D). In all cases, the power spectra comprising panels iii show mean power 4-6 min after injection, during which the increase in beta power was maximal after 30 mg/kg tianeptine administration [see Fig. 5C(iii)]. Power in the 48-52-Hz range is omitted throughout owing to the presence of 50-Hz line noise in the LFP signal.

Although the analysis presented in Fig. 4 provides a clear qualitative indication of tianeptine’s effects on LFP activity, a measure of the change in spectral power between the last vehicle baseline day and the first day of tianeptine administration (Fig 5A) provides a more appropriate measure by eliminating variability caused by differences in baseline LFP power between mice. Fig. 5B(i) shows the difference in spectral power between these two days at each frequency and time point. Note the modest increase in power with a peak at 25-30 Hz, a change that was maximal 6-8 min after injection. Fig. 5B(ii) shows the change in power as a function of frequency at this 6-8 min time-point only, with the grey area representing beta power in the 15-30 Hz range. The time course of the change in beta power is plotted in Fig. 5B(iii). Note the peak change 6-8 min after injection, and the return to baseline values within 20-30 min. This change is summarised by the area-under-the-curve (AUC) across the first 30-min of the recording session (grey shaded area), a value that is plotted in Fig. 5B(iv); a one-sample t-test revealed that this increase was significant [t(9) = 3.03; p = 0.014]. The effects of 30 mg/kg tianeptine are presented in the same way in panels 6C-E. A larger increase in beta power was evident after 30 mg/kg tianeptine administration in WT mice (Fig 5C), this time reaching a peak 4-6 min after injection and returning to baseline within 30-40 min, despite the presence of robust locomotor increases at this time point and dose (cf. Fig. 3F). A one-sample t-test of the AUC of the change in beta power over the first 30 min of the session revealed that this increase was highly significant [t(7) = 3.82; p = 0.0065; Fig. 5C(iv)]. Fig. 5D shows a similar increase and time course of beta power in δ -/- mice [increase in AUC of beta power change: t(7) = 4.01; p = 0.0051; one-sample t-test; Fig. 5D(iv)]. Fig 5E shows that, in contrast, no change in beta power was evident in µ -/- mice [increase in AUC of beta power change: t(6) = 1.04; p = 0.33; one-sample t-test; Fig. 5E(iv)]. A comparison of the AUC of the change in beta power (see Fig. 5, panels iv, for the units of this measure) after 10 mg/kg versus 30 mg/kg tianeptine in WT mice revealed a significant dose-dependent increase [10 mg/kg AUC = 2.10 ± 0.69; 30 mg/kg AUC = 7.0 ± 0.74; t(16) = 2.72; p = 0.015; independent samples t-test; Fig. 5B(iv) and 5C(iv)]. A separate analysis of the AUC of the change in beta power across the 3 genotypes (WT, δ -/-, and µ -/-) injected with 30 mg/kg tianeptine revealed a significant main effect of group [WT AUC = 7.0 ± 0.74; δ -/- AUC = 6.03 ± 1.51; µ -/- AUC = 0.20 ± 0.74; F(2,20) = 5.86; p = 0.012; ANOVA], and post-hoc pairwise comparisons (Tukey’s HSD) revealed significantly higher beta power in the WT versus µ -/-groups [p = 0.012], and δ -/- versus µ -/- groups [p = 0.032], but no difference between WT and δ -/- genotypes [p = 0.89]. This analysis confirms that tianeptine’s enhancement of beta power is dependent on the presence of µ-ORs, but does not require δ-ORs.

**Fig. 5.**
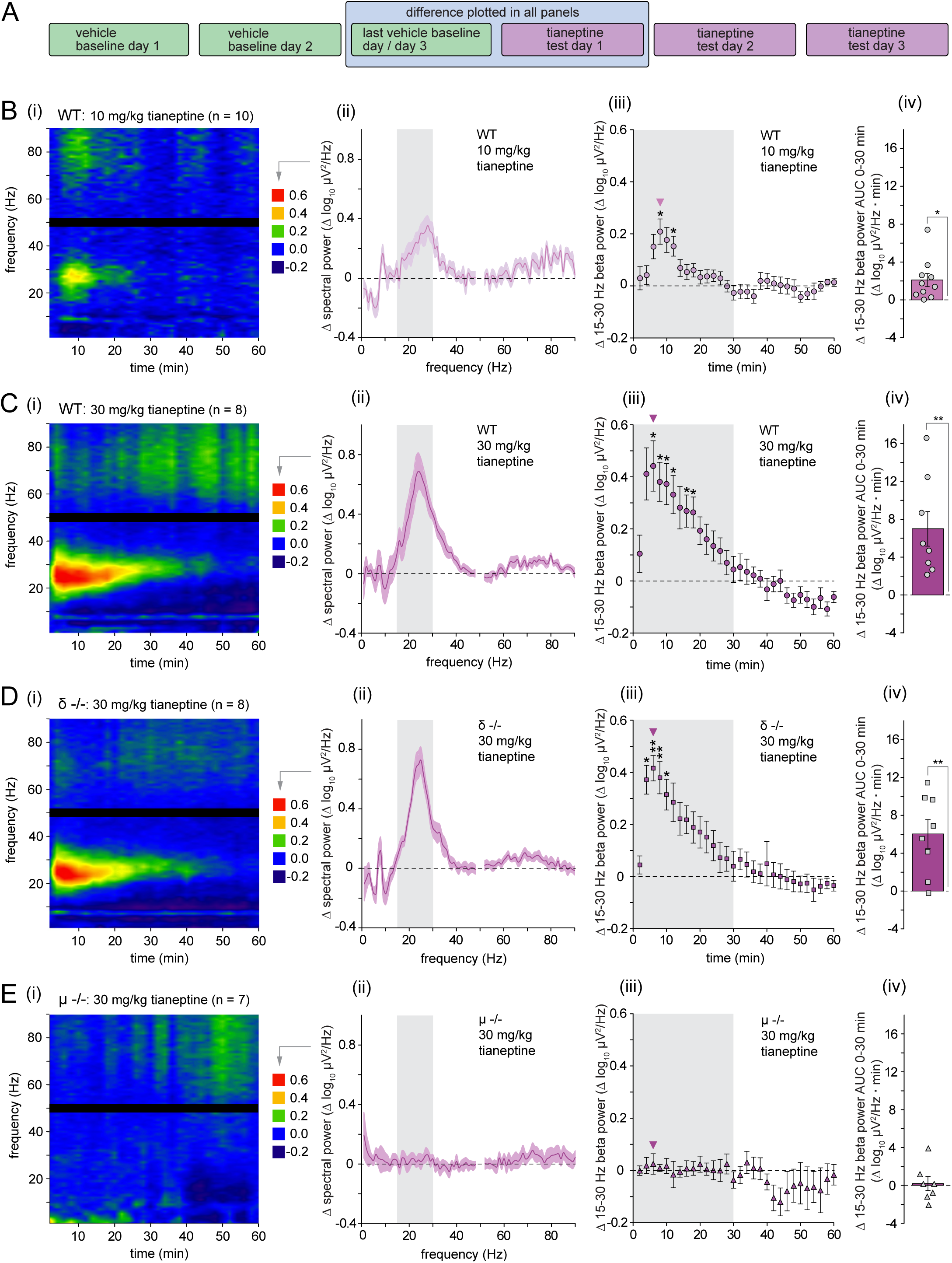
Change in spectral power after tianeptine administration. (A) Experimental timeline highlighting the focus, in this figure, in the increase in LFP power between the two days indicated by the blue rectangle (B)(i) Time-frequency heat map of the mean change in spectral power (z-axis) after 10 mg/kg tianeptine administration in WT mice (Experiment 2A), relative to the last vehicle baseline day. (B)(ii) Mean change in spectral power 6-8 min after tianeptine versus vehicle injection. (B)(iii) Mean time course of the increase in beta-frequency power (15-30 Hz; grey rectangle in panel ii after tianeptine relative to vehicle injection. Note the peak at the 6-8 min time-point. (B)(iv) Mean area under the curve (AUC) of the increase in beta power over the first 30 min after injection, corresponding to the shaded area in panel iii. Subsequent panels show data plotted in the same way for mice with wire electrode implants injected with 30 mg/kg tianeptine (experiment 2B)—WT (C), δ -/- (D), and µ -/- (E). In all cases, the mean spectral change plotted in panels ii corresponds to a 2-min period, 4-6 min after injection, during which the increase in beta power was maximal as indicated in panels iii. Power in the 48-52-Hz range is omitted throughout owing to the presence of 50-Hz line noise in the LFP signal. In panels iii: *p < 0.05; ⁑p < 0.01; one-sample t-tests with Bonferroni comparison; first 30 min of session only (grey shaded area).

### Effect of tianeptine on spatial learning in a cross-maze task

Injection of 10 mg/kg tianeptine enhanced spatial learning in a cross-maze reference memory task, but only in WT mice, not in µ or δ knockouts (Fig. 6). A photograph of the maze is shown in Fig. 6A, illustrating the presence of feeder bottles for soya milk reward in the E and W arms (bottom left and top right corners of the image). An illustration of the training procedure is provided in Fig. 6B. An individual mouse always started each training trial from the same location (either the N or S arm), but probe trials always started from the opposite location; see Materials and Methods for further details. WT mice injected with tianeptine 30 min before the start of testing showed an increase in the percentage of correct arm choices on training trials relative to vehicle-treated controls [Fig. 6C(i)]; each training block represents the average of 7 trials per day for 5 days, i.e. 35 trials. Owing to the presence of tied values, we used a non-parametric approach to the statistical analysis of percentage response data. Friedman tests revealed a highly significant improvement in percentage correct choices across trials in both vehicle [χ**^2^** = 23.6; p < 0.001] and tianeptine groups [χ**^2^** = 27.4; p < 0.001]. Mann-Whitney U-tests of performance on each trial revealed significantly higher choice performance in tianeptine-versus vehicle-treated mice during training block 3 [z = 2.84; Bonferroni corrected p = 0.018] and block 4 [z = 3.44; Bonferroni corrected p = 0.023] [Fig. 6C(i)]. To explore the use of egocentric versus allocentric strategies during learning, we adopted a stringent criterion for allocentric responding that required an allocentric response during a probe trial followed by a correct response during the subsequent training trial (Fig. 6B, inset in the lower centre of the panel; see Materials and Methods). Tianeptine-treated mice showed a sharp increase in the adoption of this strategy between training blocks 1 and 2, whereas little change was seen in vehicle-treated mice [Fig. 6C(ii)]. Mann-Whitney U-tests of stringent allocentric responding during each training block revealed significantly higher allocentric strategy use in tianeptine-versus vehicle-treated mice on block 2 [z = 2.70; Bonferroni-corrected p = 0.028] with tianeptine-treated mice showing stringent allocentric responses on 71.7 ± 10.6 % of testing days, compared to only 28.3 ± 7.16 % of controls. Fig. 6C(iii) shows the mean latencies of WT mice to choose an arm regardless of whether the choice was correct or incorrect, i.e. the time between the opening of the gate to the start box until the whole body of the mouse crossed into the E or W arm, indicated in pink in the inset illustration. An ANOVA revealed a significant decrease in escape latencies across training blocks [F(1.02, 22.4) = 11.2; p = 0.003; Greenhouse-Geisser correction], with higher latencies on the first block reflecting higher uncertainty regarding the task demands and reward location. However, there was no difference between tianeptine- and vehicle treated groups on this measure [F(1,22) = 0.25; p = 0.62]. This result is consistent with the observation that tianeptine-induced increases in locomotor activity return to near-baseline levels within ∼30 min of injection, i.e. the time point at which cross-maze testing began.

**Fig. 6.**
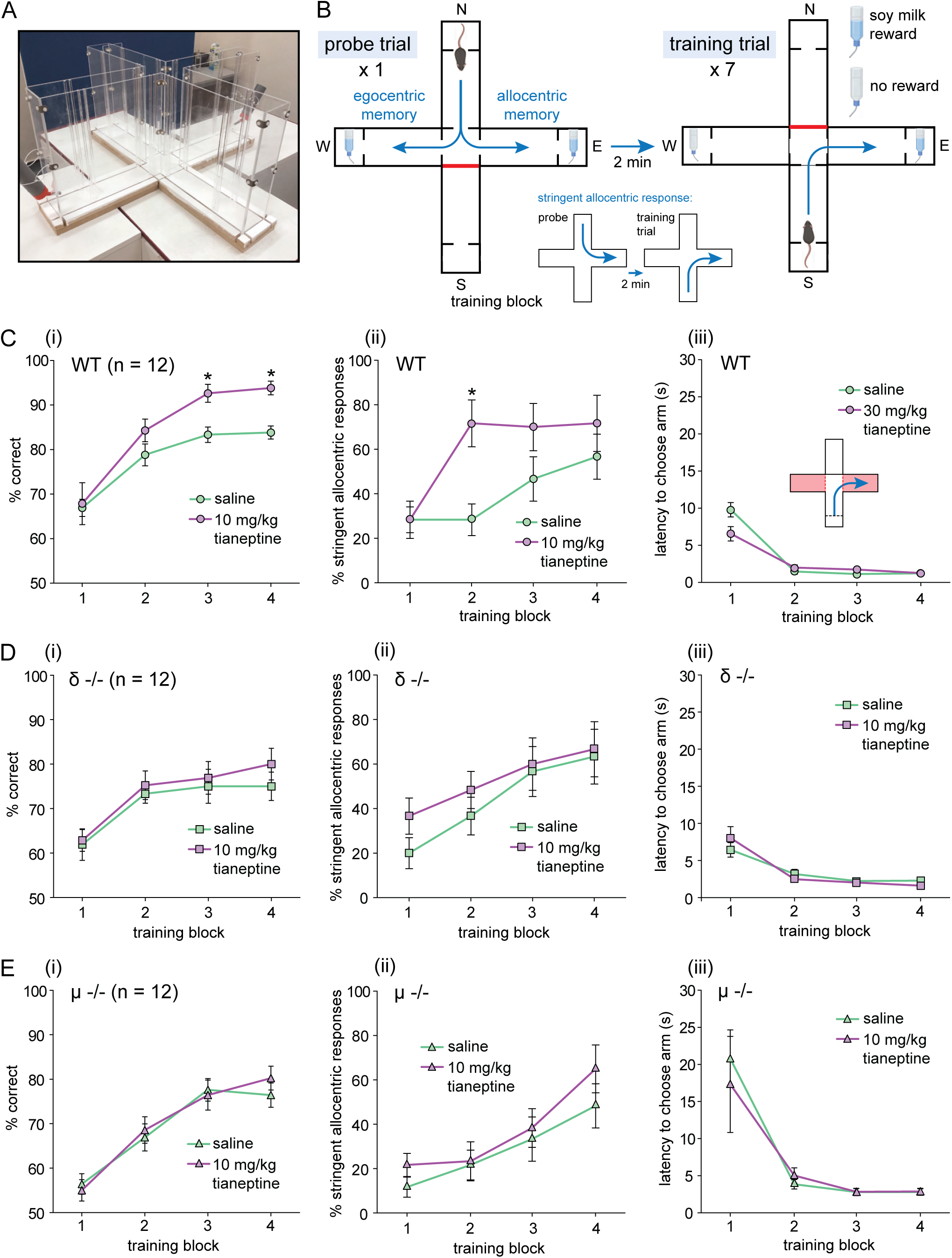
(A) Photograph of the cross-maze used in Experiment 3 (see Fig. 1). The start arms are visible in the upper left and bottom right of the image, and the goal arms with bottles to deliver soya milk reward are located in the upper right and bottom left. (B) Illustration of the apparatus and training procedure. Each testing day began with a probe trial (left-hand panel) in which mice were released from the start box opposite to their usual assigned start location in the N or S arms (N in this example). This was followed by 7 training trials starting from the usual location (S in this case). On probe trials, a response leading to the correct cardinal location (E in this case) was labelled as ‘allocentric’ whereas a turn in the direction that would have led to reward during training trials (a left turn in this example) was labelled as ‘egocentric’. A more stringent criterion for allocentric responding is provided by the occurrence of an allocentric response on the probe trial followed by a correct response on the subsequent training trial (see inset in the lower centre of panel B). To meet this criterion, a mouse must make turns in opposite egocentric directions on two consecutive trials. (C)(i) Percentage correct (i.e. rewarded) responses during training trials for vehicle and 10 mg/kg tianeptine sub-groups of WT mice. Each testing block comprises 5 days of 7 trials per day. *p < 0.05; post-hoc pairwise comparisons (Tukey’s HSD; see main text). (C)(ii) Percentage of ‘stringent allocentric’ responses made by WT mice, defined as illustrated in panel B. *p < 0.05; post-hoc pairwise comparisons (Tukey’s HSD; see main text). (C)(iii) Latency to choose a goal arm for WT mice, defined as the time between the opening of the gate to the start box and whole-body entry into the E or W arm (pink area in the inset figure). (D) Performance of δ -/- mice analysed and presented in the same way as for WT mice above. (E) Performance of µ -/- mice analysed and presented in the same way as for WT and δ -/- mice above.

In contrast to the effects of tianeptine in WT mice, the drug caused no improvement of memory in δ -/- mice (Fig. 6D). Friedman tests indicated a highly significant improvement in percentage correct choices across trials in both vehicle [χ**^2^** = 12.7; p = 0.0054] and tianeptine groups [χ**^2^** = 13.3; p = 0 0040], indicating that the mice were able to learn the task [Fig. 6D(i)]. However, Mann-Whitney U-tests with Bonferroni correction for multiple comparisons did not show differences between tianeptine and vehicle groups on any training bock, either in terms of correct choices during training trials [p > 0.9 in all cases; Fig. 6D(i)] or stringent allocentric responding [p > 0.5 in all cases; Fig. 6D(ii)]. Like WTs, δ -/- mice showed a decrease in the latency to choose an arm during training trials [F(1.31, 28.8) = 28.1; p < 0.001], but there was no difference between tianeptine and vehicle groups [F(1,22) = 0; p = 1.0].

Tianeptine also had no effect on learning in µ -/- mice (Fig. 6E). Friedman tests indicated a highly significant improvement in percentage correct choices across trials in both vehicle [χ**^2^** = 25.8; p < 0.001] and tianeptine groups [χ**^2^** = 26.5; p < 0.001], indicating successful learning [Fig. 6E(i)], but no differences between tianeptine and vehicle groups on any training bock, either in terms of correct choices during training trials [p > 0.9 in all cases; Mann-Whitney U-tests with Bonferroni correction; Fig. 6E(i)] or stringent allocentric responding [p > 0.5 in all cases; Mann-Whitney U-tests with Bonferroni correction; Fig. 6D(ii)]. Latency to choose an arm decreased during training trials [F(1.02,22.5) = 19.2; p < 0.001], but as in the other genotypes there was no difference between tianeptine and vehicle groups [F(1,22) = 0.06; p = 0.81].

## Discussion

Our central findings are that tianeptine causes a dose-dependent increase in hippocampal beta-frequency oscillations in freely moving mice, and a facilitation of spatial learning in a cross-maze reference memory task. The increase in beta power was completely blocked by deletion of the µ-OR but unaffected in δ-OR knockout animals. In contrast, the tianeptine-induced enhancement of memory was blocked in both µ -/- and δ -/- mice, implicating both receptor subtypes in the pro-cognitive actions of the drug. In parallel, we carried out a detailed analysis of the dose-dependence and time-course of the increase in locomotor activity after tianeptine administration, a phenomenon with potential implications for interpreting changes in LFP activity and learning ability, as discussed below.

An acute increase in locomotor activity is one of the hallmarks of µ-OR receptor activation in rodents. This response sometimes increases with repeated administration (Babbini and Davis, 1972), a phenomenon known as behavioural sensitisation. A similar increase in locomotor activity has previously been reported after injection of tianeptine in mice (Samuels et al., 2017), an effect that is dependent on µ-ORs located on striatal D1-receptor-expressing medium spiny neurons (Han et al., 2022). In this and previous studies, the absence of tianeptine-induced hyperlocomotion in µ knockout mice, and the lack of an effect of δ deletion suggests that activation of δ-ORs plays no role in tianeptine’s locomotor effects. Whereas some selective δ-OR agonists such as SNC80 can increase activity levels in mice, others do not, possibly owing to based signalling (Chu Sin Chung & Kieffer, 2013; Nozaki et al., 2014). In contrast to the sensitisation of the locomotor response that is sometimes observed after morphine administration, we observed no further increase in locomotor activity across three days of tianeptine injection in the current study. The lower efficacy of tianeptine relative to morphine may be responsible for this difference. In µ +/- mice with reduced receptor availability, for example, morphine still causes an acute increase in activity, but this does not increase with repeated daily injections (Bull et al., 2017a).

The main goal behind our assessment of tianeptine’s locomotor effects, however, was to characterise a ‘side effect’ of the drug with potential implications for the interpretation of its effects on hippocampal network activity and spatial learning. Regarding the first of these actions, we have previously reported that tianeptine causes an increase in hippocampal beta-frequency oscillations in anaesthetised rats. Our current data extend these findings to awake mice, and establish that the phenomenon is dependent on the presence of µ- but not δ-ORs. Importantly, the time-course of this effect was dissociable from tianeptine’s locomotor effects; at a dose of 30 mg/kg, beta power reached a peak ∼6 min after injection, in contrast to the increase in locomotor activity that peaked >20 min after injection, and remained elevated long after beta power had returned to baseline. In this regard, the time course of tianeptine’s LFP effects arguably best parallels the pharmacokinetics of tianeptine itself, rather than its metabolite MC5 (cf. Samuels et al., 2017). For several reasons, it is unlikely that the enhancement of beta activity is solely an artifact of tianeptine-induced hyperlocomotion: first, the time courses of the two phenomena are distinct; second, activity in the theta and high-gamma ranges is more strongly associated with increased activity and running speed (Whishaw & Vanderwolf, 1973; Chen et al., 2011; Ahmed & Mehta, 2013) and a relatively selective increase in beta power is not consistent with these observations; and third, increases in beta power are observed even in the absence of locomotion under anaesthesia (Burt et al., 2026).

Discussion of the functional significance of tianeptine’s enhancement of beta oscillations is necessarily speculative at this stage. Cortical beta oscillations have long been associated with sensorimotor processes (Barone & Rossiter 2021), but oscillations within this frequency range are likely to play a range of distinct roles in different brain networks, including cognitive operations (Lundquist et al., 2024). Coherent beta-frequency cortico-hippocampal oscillations are, for example, observed during the learning of sensory discriminations, and beta also been implicated in the coupling of hippocampus-dependent memory with brain areas such as the prefrontal cortex and ventral striatum that mediate decision-making, action selection, and reward (Miles et al., 2023). Conversely, reduced EEG beta power is observed in Alzheimer’s (Kopcanova et al., 2024). Hippocampal beta oscillations occur in rodents during the exploration of novel objects and environments (Berke, 2008; França et al., 2014), and their power predicts subsequent memory recall (Iwasaki et al., 2021). In view of these observations, it is possible that a pharmacological enhancement of endogenous beta activity might facilitate memory formation. Enhancing endogenous beta oscillations, or driving rhythmic activity at this frequency, might also have therapeutic value in depression, for example by shifting activity away from ruminative activity associated with activation of the default mode network (Chou et al., 2023), and towards the processing of sensory information and goal-directed action (cf. Miles et al., 2023).

Consistent with this view, changes in beta and gamma power have been implicated in the pathophysiology of depression (Fitzgerald & Watson 2018; Yin & Li, 2026) and tianeptine is not alone among antidepressants in causing marked increases in oscillatory activity in these frequency bands—ketamine, for example, is well known to increase cortical and hippocampal gamma power (Gilbert & Zarate Jr, 2020). Unlike ketamine and tianeptine (see Burt et al., 2026), SSRIs do not acutely enhance beta or gamma activity in rodents, but chronic administration results in increases (Eskikurt et al., 2024), and elevated cortical beta is predictive of antidepressant response in humans (Papp et al., 2018; Li et al., 2026). However, it is unlikely that beta oscillations provide a specific biomarker for antidepressant efficacy since several classes of positive allosteric modulators of the GABAA receptor such as neurosteroids and benzodiazepines cause increases in hippocampal beta oscillations, but whereas both classes of drug have anxiolytic actions, only the former is effective in depression (Lim et al., 2020; Lambert et al., 2023; Takasu et al., 2024).

The mechanisms by which tianeptine increases hippocampal beta activity remains unclear, but other µ-OR agonists such as buprenorphine and morphine can also increase power in the beta and gamma ranges (Reakkamnuan et al., 2015; Sakae & Martin, 2019; Martin et al., 2025; Burt et al., 2026). Hippocampal µ-ORs are heavily expressed in interneurons, particularly those mediating perisomatic sites (e.g. Stumm et al., 2004). Analogous to their actions in the mesolimbic dopamine system (e.g. Bull et al., 2017b), agonists cause a reduction in GABA release and / or a reduction in firing rate, leading to a disinhibition of principal neurons (Zieglgänsberger et al., 1979; Neumaier et al., 1988; Lambert et al., 1991; McQuiston & Saggau, 2003; Shao et al., 2020). Consistent with this action, we have previously observed that tianeptine causes a reduction in the frequency of spontaneous inhibitory postsynaptic currents (IPSCs) and a decrease in the amplitude of evoked IPSPs in CA1 pyramidal neurons in vitro; tianeptine also causes a reduction in paired-pulse inhibition of the CA1 population spike, a measure of the strength of inhibitory transmission, in vivo (Martin et al., 2025). Nonetheless, pharmacological disinhibition by local antagonism of GABA_A_ receptors does not enhance power at the frequencies enhanced by tianeptine in the current study (Martin et al., 2025), though an increase in power at lower frequencies has been observed (Gwilt et al., 2020). Similarly, reducing the activity of perisomatic interneurons such as PV+ cells does not enhance beta oscillations, but instead reduces gamma power (Fuchs et al., 2007; Sohal., 2012; Antonoudiou et al., 2020). These findings suggest that additional or alternative mechanisms are likely to be involved in tianeptine’s actions. A possibility is the ability of tianeptine to increase AMPA-receptor-mediated synaptic transmission by phosphorylating GluA1 subunits (Svenningsson et al., 2007; Szegedi et al., 2011) and enhancing evoked excitatory postsynaptic currents (eEPSCs), a phenomenon that we have recently observed in CA1 pyramidal neurons (Mariano et al., 2026).

The dual ability of tianeptine to disinhibit excitatory neurons and enhance AMPA-receptor-mediated excitatory drive might underlie its ability to enhance the formation of long-term potentiation (LTP) of synaptic strength and hippocampus-dependent memory. Tianeptine’s memory-enhancing actions have been well documented, particularly its ability to ameliorate impairments caused by stress or neuropathology (Zoladz et al., 2008). However, most of these studies predate knowledge of tianeptine’s interactions with opioid receptors, and the receptors responsible for its pro-cognitive effects were unknown. In the present study, we find that tianeptine improves spatial learning in experimentally naïve wild-type mice, and that this improvement is absent in mice lacking either the µ- or δ-OR, implicating both subtypes in the drug’s pro-cognitive effects. A role for δ-ORs is supported by previous evidence implicating hippocampal δ-ORs in hippocampus-dependent learning, including the formation of allocentric spatial memory (Le Merrer et al., 2013; Pellissier et al., 2016). The involvement of hippocampal µ-ORs in learning and memory processes is perhaps more surprising, considering that systemic administration of µ-ORs often impairs, rather than facilitates performance in spatial learning tasks in rats and mice (e.g. McNamara & Skelton, 1991; Zhu et al., 2011). However, the low efficacy of tianeptine relative to canonical agonists such as morphine might be relevant here. The intense reward, hyperlocomotion, and motivational changes associated with systemic administration of opioid receptor agonists may mask potential memory-enhancing effects (cf. McNamara & Skelton, 1991) that might be evident only at very low doses, or following intracerebral administration. Note that cross-maze training in this study was carried out at a dose and time point that did not result in hyperlocomotion at the time of memory formation—although this observation raises additional unanswered questions concerning the pharmacodynamics of tianeptine’s actions and those of its metabolite MC5, including the possibility of persistent effects on downstream signalling pathways. Our findings also raise questions concerning the physiological role of endogenous opioid signalling in learning and memory. Knockout of µ-ORs can impair spatial learning under some circumstances, suggesting a role for endogenous µ signalling in memory formation (Jamot et al., 2003; McQuiston & Saggau 2003), although stress-induced activation of µ-ORs can, conversely, impair the retrieval of existing memory (Shi et al. 2020). Endogenous opioids can also be released in the hippocampus in response to novelty (Netto, 2022), suggesting that opioid receptors might be members of a wider group of G-protein-coupled receptors involved in the novelty-induced modulation of memory by, including dopamine D1 receptors (Tse et al., 2023) and metabotropic glutamate receptors (Gravius et al., 2010).

The site and circuit-level mechanisms of tianeptine’s pro-cognitive actions are also unclear although the hippocampus is a likely target, and hippocampal activity is required for allocentric learning in a task similar to the one used in the current study (Packard & McGaugh, 1996). The hippocampus of both humans and rodents shows gradients of anatomical connectivity and functional specialisation along its long axis (Strange et al., 2014)—with the dorsal-ventral axis in rodents being homologous to the posterior-anterior axis in humans. The division of labour along this axis is graded rather than absolute (de Hoz & Martin, 2014), but the dorsal / posterior hippocampus is more strongly associated with allocentric spatial learning than the ventral / anterior hippocampus that plays a larger role in motivation, emotion, and reward-related processes owing to its connectivity with structures such as the ventral striatum and prefrontal cortex (Moser & Moser, 1998; Bannerman et al., 2004). The memory-enhancing and antidepressant actions of tianeptine might therefore reflect a common action along the dorsoventral axis of the hippocampus. Our observation of the sharp increase in the rate of adoption of an allocentric strategy in tianeptine-treated WT mice is consistent with the activation of dorsal hippocampal opioid receptors. In contrast, existing evidence suggests that the activation of ventral hippocampal µ-ORs plays a larger role in tianeptine’s antidepressant actions (Han et al., 2022). Since memory impairment is a common feature of depressive illness, and depression is common in people with neurodegenerative disorders (Yang et al., 2026), this dual action is a highly desirable feature in an antidepressant drug.

While the example of tianeptine illustrates the therapeutic potential of opioid receptor activation for cognitive enhancement and the treatment of depression, the rewarding and addictive properties of opioid receptor agonists limit the clinical utility of this class of drug (see Edinoff et al., 2023). Further research into the circuit-level roles of hippocampal µ- and δ-ORs, and their downstream signalling pathways in specific cell types, might provide a framework for novel therapeutic approaches that avoid these drawbacks. For example, we have recently identified a novel signalling mechanism linking the activation of postsynaptic µ- and δ-ORs to the enhancement of synaptic currents mediated by calcium-permeable AMPA receptors (Mariano et al., 2026), a mechanism that might be central to the enhancement of memory that depends on the activation of these receptors. Owing to its favourable side-effect profile, δ-OR signalling presents an appealing target for future work on cognitive enhancement whereas µ-OR signalling provides a potent mechanism for driving hippocampal beta oscillations, one that is likely to be dissociable from the mesolimbic mechanisms of drug reward.

## Acknowledgements

This work was funded by an MRC Doctoral Training Programme studentship held by Margarida Trigo. We would like to thank Christian Keitel for providing Python code used for preprocessing of electrophysiological data, Patrick Spooner for design and 3-D printing of the commutator assembly, and Biological Services staff for animal care and husbandry.

